# Analysis of mixtures of birds and insects in weather radar data

**DOI:** 10.1101/2024.07.17.601450

**Authors:** Xu Shi, Jacob Drucker, Jason W. Chapman, Melissa Sanchez Herrera, Adriaan M. Dokter

## Abstract

Weather radars are increasingly used to study the spatial-temporal dynamics of airborne birds and insects. These two taxa often co-occur and separating their contributions remains a major analytical challenge. Most studies have restricted analyses to locations, seasons, and periods when one or the other taxa dominates. In this study, we describe an analytical method to estimate the proportion of birds and insects from vertical profiles of biological reflectivities, using a minimal number of assumptions on the airspeeds of birds and insects. We evaluated our method on understudied regions where airborne insect density is too high for existing approaches of studying bird migration with weather radars: the tropics (Colombia) and the southern temperate zone (Southeast Australia). Our method estimates that bird and insect signals routinely reach similar magnitudes in these regions. Retrieved patterns across daily and annual cycles reflected expected biological patterns that are indicative of migratory and non-migratory movements in both climates and migration systems. Compared to fixed airspeed thresholding, we obtain finer separation and retain more spatial-temporal complexity that is crucial to revealing aerial habitat use of both taxa. Our analytical procedure is readily implemented into existing software, empowering ecologists to explore aerial ecosystems outside the northern temperate zone, as well as diurnal migration of birds and insects that remains heavily understudied.

**Lay summary:**

- We developed a new method to differentiate between birds and insects in weather radar data.
- This method uses minimal assumptions about the flight speeds of birds and insects.
- We tested the method in regions with high insect density: the tropics (Colombia) and southern temperate zone (Southeast Australia).
- Our method estimated proportions of birds and insects that captured expected patterns of daily and annual movements, which were indicative of migratory and non-migratory movement of both taxa.
- Unlike fixed airspeed criteria for bird and insect separation, our approach provides a more detailed understanding of aerial habitat use by both birds and insects.
- This method can be easily added to existing software, helping ecologists study bird and insect movements in less-studied areas and ecosystems.

## Introduction

Weather radars have been increasingly used to quantify and study the spatial-temporal dynamics of airborne birds and insects (Bauer et al. 2017). Both taxa can be incredibly numerous in the air and constitute the main biological scattering on radar, especially during migration when populations are most mobile (Hu et al. 2016, Dokter et al. 2018, Wotton et al. 2019, Nussbaumer et al. 2021b, Huang et al. 2024). Even though the radar cross-section of insects is often orders of magnitudes smaller than for birds, insects can be present in enormous numbers, such that bird and insect scattering can reach similar reflectivities. (Leskinen et al. 2011, Westbrook and Eyster 2017, Cui et al. 2019, Stepanian et al. 2020). Distinction between these two main taxa in radar products is thus imperative for reliable interpretation of their airborne movements (Bauer et al. 2019). To date, almost all bird migration studies using weather radar are carried out in the northern temperate zone during the migratory periods of birds. These studies have been able to mostly side-step the problem of separating insect and bird signals co-occurring at similar magnitudes. In temperate latitudes, biological scattering is heavily dominated by birds during the peak migration season at night, while insects become more numerous during the day (Rennie 2014, Hu et al. 2024). Consequently, diurnal studies of birds have been challenging, resulting in a focus on using nocturnal data in bird studies (Van Doren and Horton 2018, Dokter et al. 2018, Nilsson et al. 2019), and limited weather radar-based study of insects.

However, assuming a diel divide between airborne birds and insects is a clear oversimplification of biological reality. Diurnal migrating birds often fly at the same time as insects that take to the air for various purposes besides migration (Chapman et al. 2011), while insect migration can be extremely intense at night (Westbrook and Eyster 2017, Huang et al. 2024, Haest et al. 2024). In temperate regions during early fall and late spring migration, bird numbers are low while insects can become very abundant, their reflectivity values can become comparable (Leskinen et al. 2011, Westbrook and Eyster 2017, Nussbaumer et al. 2021a). Moreover, our understanding of the timing and volume of bird and insect fluxes are heavily biased towards the northern temperate zone. Whether insect or bird movements dominate biological signals during certain seasonal and diurnal periods has yet to be evaluated, especially in the tropics and in the Southern Hemisphere. These scenarios require dedicated methods to distinguish between birds and insects when they are aloft simultaneously and co-occurring at similar reflectivities.

Currently, filtering insects from bird signals in weather radar products is mostly based on the difference in their velocity properties. Several studies used a fixed self-powered airspeed threshold (e.g. 5 m/s) to reduce contamination from the usually slow-flying insects (Larkin 1991, Horton et al. 2016b, 2018). However, strict thresholds may be crude given overlapping airspeeds between large, fast-flying insects and small, slow-flying birds, particularly under variable wind assistance (Cabrera-Cruz et al. 2013). Migrating birds also show a greater spatial variability in velocity than wind-borne hydrometeors and insects, as measured through standard deviations (texture) of radial velocity (Gasteren et al. 2008, Dokter et al. 2011). Removing altitude layers with a radial velocity standard deviation (sd_vvp) below 2 m/s showed satisfactory results in removing insect-dominated cases in C-band radars that operate in dual pulse repetition frequency mode. Variability in velocity has not been thoroughly examined in S-band radars (Dokter et al. 2011, 2019), or when birds and insects co-occur in mixtures of comparable magnitude.

In specific geographic regions, bimodal distributions can be observed in both airspeed and radial velocity standard deviation (sd_vvp), with the relative magnitude of insect and bird components in those full distributions changing throughout the season. Nussbaumer et al. (2021b) analyzed these patterns using gaussian mixture modeling in western Europe. These authors compared measured airspeed and sd_vvp values to bimodal airspeed and sd_vvp distributions obtained over longer time periods to infer proportions of birds and insects. We have found that bimodal distributions of airspeed are not always present, especially in climates where insects are common year-round, therefore the robustness and general applicability of this decomposition approach has limitations.

Dual-polarization radar measurements can also assist in the differentiation between bird and insect scatters. Insects tend to exhibit high differential reflectivity values due to their elongated body shapes (Stepanian et al. 2016), but large overlap exists with birds and dual-polarization metrics are unavailable in global repositories of single-polarization data.

In summary, there is a continued need for a general approach for estimating taxonomic proportions across varied ecosystem scenarios, radar systems and data types. In this study, we propose a simple analytical method to estimate the proportion of birds and insects from vertical profiles of biological reflectivities, which relies only on a minimum number of assumptions on the self-powered airspeeds of birds and insects. We propose to use prior information on the self-powered airspeeds of birds and insects in analyses, and to use these assumptions to separate bird and insect contributions using analytical expressions that can be applied readily to vertical profile data. To test our approach and explore the bird-insect proportions in climates other than the northern temperate zone, we applied this method on data from tropical climates (Colombia) and southern temperate zones (Southeast Australia), where insect contributions are expected to be dramatically higher than in northern temperate regions due to a lower magnitude of bird migration, higher insect abundance, and greater variation in insect body size compared to the temperate northern hemisphere.

We tested the above equations on vertical profiles generated by the vol2bird algorithm (Dokter et al. 2019) on Australian and Colombian weather radar data. We hypothesized that our model estimates of migrating bird and insect proportions would reflect known biological patterns in case studies across daily and annual cycles. For example, we would expect our model to estimate higher proportions of birds migrating at night and early morning, and insects to occur during afternoons as is known from other studies (Chapman et al. 2011, Drake and Reynolds 2012). Similarly, we would expect to see higher estimated proportions of birds during spring and fall when birds are migrating, and higher estimated proportions of insects during warmer or wetter seasons that are associated with reproduction and emergence (Dokter et al. 2011, Drake and Reynolds 2012).

### Radar scattering theory for birds and insects

The standard reflectivity product of weather radars is the reflectivity factor Z, which is related to the linear reflectivity η by

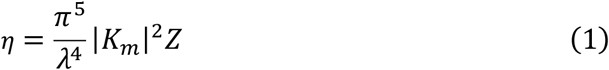

with η in aeroecology conventionally expressed in units cm^2^/km^3^ with λ the radar wavelength and |K_m_|^2^ the squared norm of the complex refractive index of water (0.93 at both S-band and C-band wavelengths, Doviak, 2006). For a mixture of birds and insects, our goal is to partition η, such that a proportion *f* should be attributed to insects, and the remaining part (1 − *f*) to birds, with 0 <= *f* <= 1. The reflectivity can then be converted into animal densities using assumptions on the effective radar cross section for each taxon:

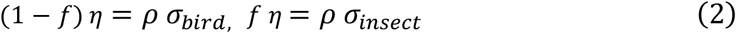

with ρ the animal density and σ the radar cross section (Dokter et al. 2011, Chilson et al. 2012). In the context of vertical profile estimation, any azimuthal dependencies in the radar cross sections are averaged out, and therefore σ is typically expressed as a single value.

The radar cross section of an animal strongly depends on its size relative to the radar wavelength λ. Insects are typically much smaller than the radar wavelength, and insect scattering falls therefore into the same Rayleigh scattering regime as precipitation, where for water droplets of diameter D the cross section can be expressed as:

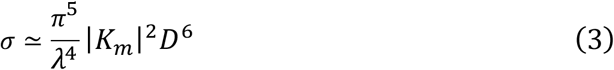

For insects we expect the same general scaling relationships with size and radar wavelength as for precipitation, i.e. a quadratic relationship with insect mass, and an inverse λ^4^ relationship with radar wavelength:

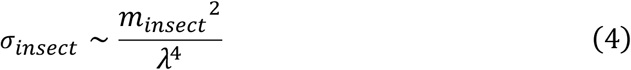

Birds are typically of similar dimension or larger than the radar wavelength, and qualitatively different scattering regimes apply to bird and insect taxa. Bird scattering occurs in the Mie scattering regime, which is the resonant transition zone towards normal scattering. Here the (azimuthally averaged) radar cross section becomes proportional to the radiated surface area of an object, i.e. proportional to the bird’s mass raised to the power 2/3:

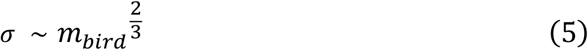

and no longer showing a strong radar wavelength dependence (cross-sections for birds can still be somewhat wavelength dependent through wavelength dependence of their refractive index). Measurements of species-specific radar cross sections indeed reveal consistent gradual increases of radar cross section with bird mass (Horton et al. 2019).

## Methods

### Bird and insect partitioning

We will use vector notation throughout, with vectors notated in bold font. We define the wind vector ***w***, the ground speed vectors ***g***_*bird*_, ***g***_*insect*_, ***g***_*mix*_ and the airspeed vectors ***a***_*bird*_, ***a***_*insect*_, ***a***_*mix*_ for birds, insects and the mixture of birds and insects, respectively. The vector’s scalar speed is given by the L2 norm of each vector, which we will notate in normal font, e.g. *g*_*bird*_=∥***g***_*bird*_∥.

The airspeed of the mixture is the vector sum of bird and insect airspeed, weighted by their relative proportions. Our goal is to express *f* - the reflectivity proportion associated with insects - in terms of the birds’ airspeed *a*_*bird*_ and the insects’ airspeed *a*_*insect*_, assuming insects move in the same direction as wind. While insects can maintain self orientation and compensate for wind drift, the angular differences between their heading and wind direction are usually small (Chapman et al. 2008, 2010) and weather radars have a much better ability to detect small insects, which are far more common and more prone to full drift (Chapman et al. 2011, Hu et al. 2016).

Winds, ground speeds and airspeeds are related through triangles of velocity, as in ***g***_*i*_= ***w*** + ***a***_*i*_ with i one of {birds, insects, mix}. Directional angles are expressed clockwise from north in radians. The mixture’s airspeed is the vector sum of the bird airspeed ***a***_*bird*_ and insect airspeed ***a***_*insect*_, where *u* and *v* are the vectoral components of the birds’ airspeed. Since insects are assumed to move in the same direction as the wind, we may write:

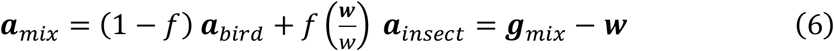

where in the last step we invoked the triangle of velocity for the mixture. Rearranging Equation 6 allows us to express the bird airspeed in terms of the mixture ground speed and the wind:

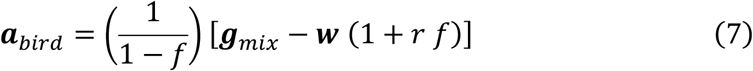

where we defined *r* as the ratio of the insect air speed to the wind speed, as in 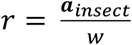. This equation shows that when insects follow the wind with zero airspeed, we have ***a***_*insect*_= *r* = 0, and therefore the bird airspeed vector becomes proportional to ***g***_*mix*_− ***w***, irrespective of the value of *f*. This also shows that the birds heading 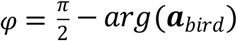 in this case does not depend on assumptions of the magnitude of the bird airspeed ***a***_*bird*_ or the proportion of insects *f*, and should therefore be a relatively robust metric whenever insect airspeeds can be assumed to be close to zero.

Calculating the norm of the bird airspeed gives us a quadratic equation for *f* that can be solved using standard algebra:

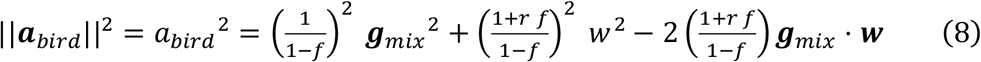

with · indicating the dot product.

Rearranging (8) to a standard quadratic equation for *f*:

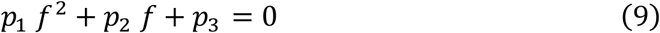

we find the following quadratic coefficients:

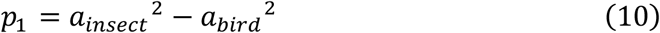

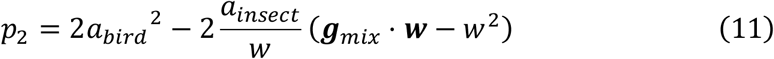

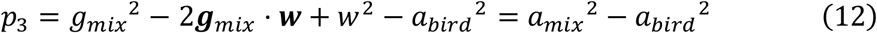

where in the last equality for *p*_3_ we invoked the law of cosines for the mixture’s triangle of velocity (see Fig.1).

**Figure 1:**
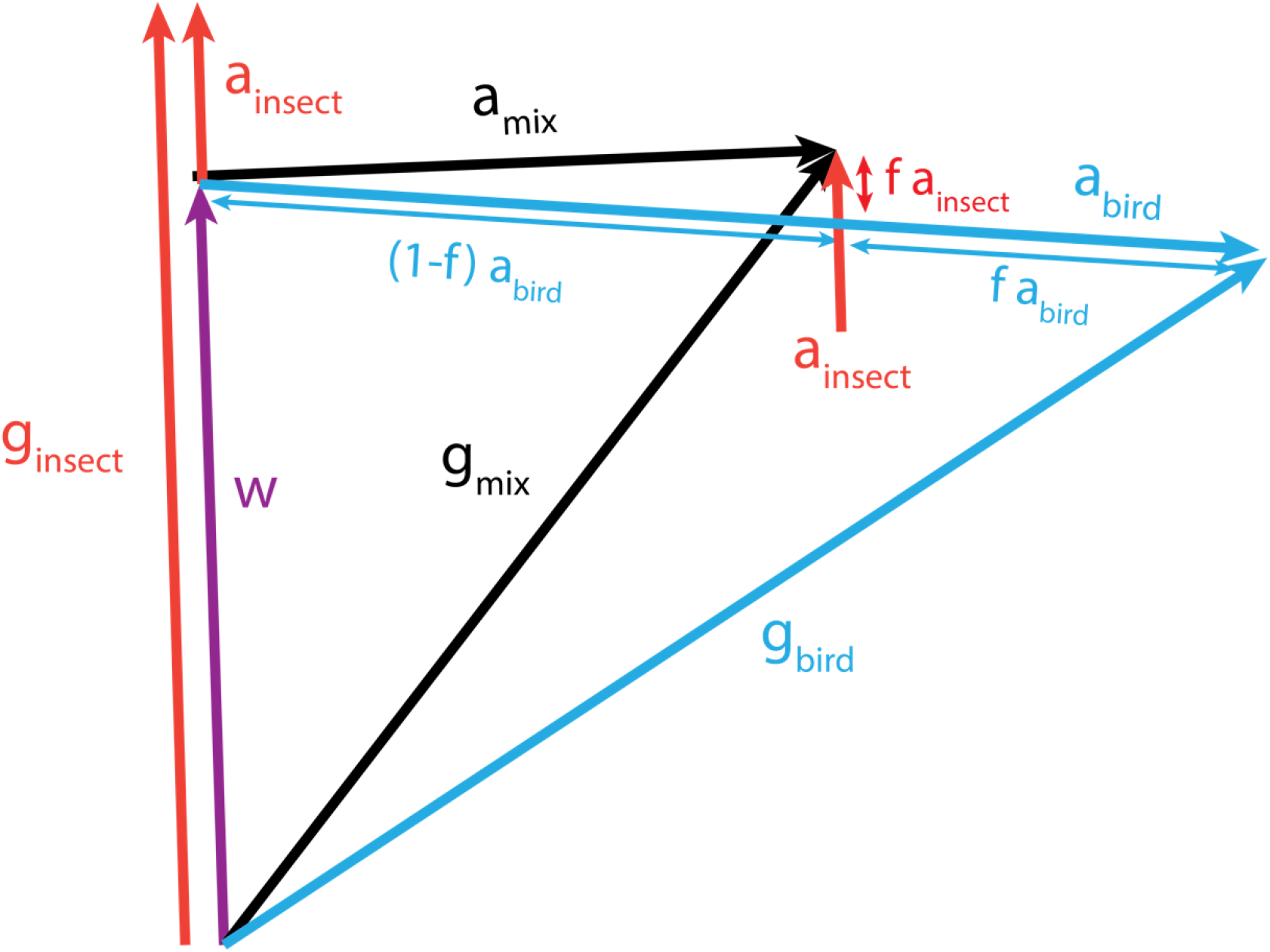
Triangles of velocities for wind vector (***w*** in purple) and the air and ground speed for the mixture (***a***_*mix*_, ***g***_*mix*_ in black), birds (***a***_*bird*_, ***g***_*bird*_ in blue) and insects (***a***_*insect*_, ***g***_*insect*_ in red), respectively.

The quadratic formula can then be used to solve *f*:

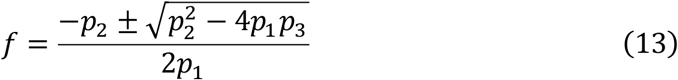

under meaningful ecological situations, we would have *a*_*bird*_ > *a*_*mix*_ > *a*_*insect*_. Applying this condition, we can show that only the solution:

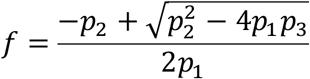

leads to positive real solutions for *f* (see supplemental methods). In cases where the measured airspeed is smaller than insect airspeed *a*_*insect*_, we assume *f* to be 1 (no birds in the mixture), while in the cases of measured airspeed being larger than bird airspeed *a, f* is assumed to be 0 (no insects in the mixture).

When the insects have no own self-speed (*a*_*insect*_= 0) we find

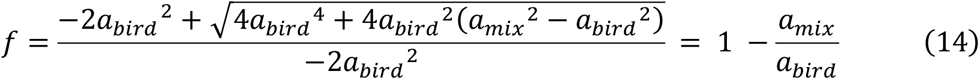

Combing the above equations, we showed that *f* and the bird airspeed vector ***a***_*bird*_ can be solved with only two parameters: the magnitude of birds’ airspeed *a*_*bird*_ and the magnitude of insects’ airspeed *a*_*insect*_. The other 4 parameters, the ground speed components ***g***_*mix*_ measured by the radar and the wind speed components ***w*** that can be obtained from a weather model or balloon sounding. We attached an R function in the supplementary methods that can be readily applied to solve *f*, and the birds’ ground and air speeds with these parameters.

### Case study sites and weather radar data

We obtained Colombian weather radar data from the country’s Institute for Hydrological, Meteorological, and Environmental Studies (IDEAM), focusing on the stations in San Jose del Guaviare, Guaviare department, south-central Colombia (name: GUA, 2.53°N, longitude: 72.62°W, elevation: 218m) and Barrancabermeja, Santander department, north-central Colombia (name: BAR, latitude: 6.93°N, longitude: 73.76°W, elevation 80m, Fig. 2A). Both of these radars are dual-polarization and operate at C-band (5.3 cm) with a beamwidth of 1°. For the Australian case study, we obtained the level-1 dataset from the Australian National Computational Infrastructure (NCI, Soderholm et al., 2019). We used data from one radar in Captains Flat, New South Wales, southeast Australia (name: CapFlat, 35.66°S, longitude: 149.51°E, elevation: 1382m, Fig. 2B) for the year 2018. This is an S-band (10.4 cm wavelength) single-polarization radar with a beamwidth of 1.9°. Reflectivity data in the Australian dataset was calibrated by matching collocated information with the Tropical Rainfall Measuring Mission (TRMM) and Global Precipitation Measurement (GPM) satellite passes (Louf and Protat 2023) before further processing.

**Figure 2:**
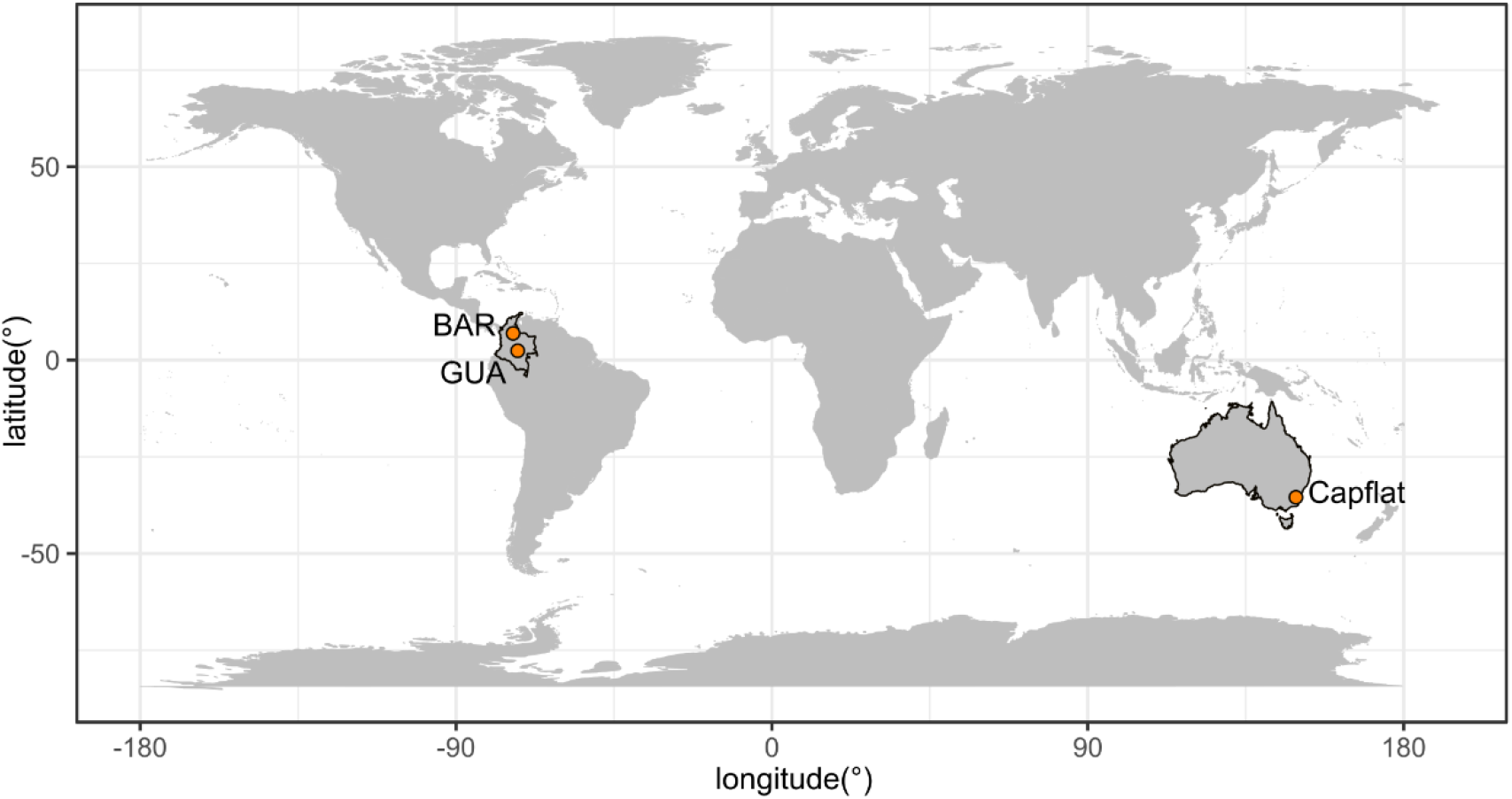
Geographical location of the weather radars included in this study. The BAR and GUA radars are located in Colombia in the tropics and (B) the CapFlat radar is located in southeast Australia in the south temperate zone. The extent of Colombia and Australia are highlighted with darker lines.

The Colombian radars are located in hot tropical lowlands, where seasonality is defined more by rainfall than temperature. The Barrancabermeja radar lies within the Magdalena River Valley but its beams reach the slopes of the Andes. San Jose del Guaviare sits at the junction between the Amazon and Orinoco basins. We chose these Colombian radars based on their low elevations that maximize the volume of airspace captured by a single scan and the continuity of the available time series. The CapFlat radar is the highest radar in Australia in terms of elevation, with winter temperature (average low in July) reaching around 1.1°C and summer temperature (average high in January) reaching around 27.9°C. Combined with its southerly latitude, it is most likely to record seasonal patterns in biological activity aloft in the temperate regions of Eastern Australia.

We selected these radars because their locations present the opportunity to test our model in both southern temperate and tropical systems. Migration in southern temperate regions often has lower volumes of bird migration relative to north temperate regions such that birds may compose a lower proportion of aerial biomass in weather radar data. Similarly, northern South America also has a lower relative abundance of migratory birds than North America, and a high abundance of insects. These sites provide a unique opportunity to study the daily and annual patterns of local biological activity in the air and the seasonal migration of the long-distance migrants from the Northern Hemisphere into and out of the neotropics. The inclusion of these contrasting environments allows a comprehensive examination of the model’s performance across a spectrum of climatic conditions.

### Calculating vertical profiles

We extracted vertical profiles of biological reflectivity (η, cm^2^·km^-3^) using the vol2bird algorithm from R package “bioRad” (Dokter et al. 2019) to temporal and altitudinal resolution of 0.1 hour and 200 m in the Australian dataset and 0.17 hours and 100 m in the Colombian dataset. Precipitation in the Australian dataset was removed using MistNet, a convolutional neural network and screened manually by visually checking time-altitude plots of reflectivity factor and removing time-altitude ranges with meteorological contamination (Lin et al. 2019, Dokter et al. 2019). Precipitation in the Colombian dataset was removed using the correlation coefficient (ρ_HV_, ρ_HV_ > 0.95 are removed, Stepanian et al., 2016). The vertical profile includes the reflectivity and the associated U (east-west) and V (south-north) components of ground speed. We retrieved The U and V components of wind speed from the ERA5 reanalysis for Australia (Hersbach et al. 2023) and the North American Regional Reanalysis for Colombia (Mesinger et al. 2006). We downloaded the hourly (Australia) to 3-hourly (Colombia) data at pressure levels from 1000 to 750 hPa, spatially interpolated to the radar location, and further interpolated to the altitudinal and temporal resolution of the vertical profiles. To summarize daily and seasonal patterns, we calculated the mean bird proportion, bird density ((1 − *f*) ∗*η*) and insect density (*f*∗*η*) for each day of the year across a data set of four years from the BAR radar in Colombia (*n =* 171,986 scans), and one year at the CapFlat radar in Australia (*n =* 105,120 scans). We chose the BAR radar because time series from this radar is more complete, and higher quality with better velocity information. To illustrate the seasonal patterns, we created yearly trends of bird proportions, bird and insect density by fitting Generalized Additive Models (GAM, Wood, 2017) with quasi-Poisson family and cyclic cubic regression splines, using bird proportions and the densities as response and day of year as predictor. Additionally, we calculated the reflectivity weighted daily mean north-ward component of *g*_*bird*_ by summing the north-ward component of ***w*** and ***a***_*bird*_ to demonstrate the seasonal variation in the north-south component of bird flight direction. We further evaluated the model’s performance by comparing it to insect filtering based on a fixed airspeed criterion of 5 m/s (Larkin 1991, Gauthreaux and Belser 1998), by comparing the total amount of bird density and the proportion of time-altitudinal bins retained throughout the study periods. We also reported two case studies with the patterns of bird proportion throughout exemplar 24-hour periods during typical migration season in CapFlat, Australia in March 2018, and San Jose del Guaviare, Colombia in April 2019 to demonstrate the model’s capacity to reveal fine scale patterns for both taxa.

### Sensitivity Analysis

Estimates of bird and insect proportions require parametrizations of the bird airspeed *a*_*bird*_ and insect airspeed *a*_*insect*_. we used a = 10 m/s as the average airspeed for birds, collating estimates from weather and tracking radar studies (Alerstam et al. 2007, Nilsson et al. 2014, Horton et al. 2016a, Nussbaumer et al. 2022). We used ***a***_*insect*_= 1 m/s to represent the whole insect community, ranging from the largest (but comparatively scarce) migrants (such as noctuid and sphingid moths, nymphalid butterflies, and dragonflies) that have self-powered airspeeds of 4-5 m/s (Chapman et al. 2010, Hedlund et al. 2021, Hu et al. 2021, Menz et al. 2022), to the vastly more numerous micro-insects with airspeeds <1 m/s that largely drift with the wind (Rennie 2014, Hu et al. 2016, Nussbaumer et al. 2021a). We further demonstrated the influence of different bird and insect airspeed values on the resulting bird proportion. We applied bird airspeed from 8 to 15 m/s and insect airspeed from 0 to 5m/s and reported the mean bird proportions retrieved for both Australian and Colombian datasets.

## Results

### Seasonal Patterns

In the absence of a ground truth on the true insect and bird abundances in our study areas, we rely on qualitative assessments of whether recovered patterns are ecologically meaningful and consistent with prior natural history knowledge. Both in Colombia and Australia, we recover distinct seasonal patterns in aerial bird and insect abundance that are consistent with large-scale seasonal migration. We find distinct seasonal patterns in the predicted proportions and abundances of nocturnal birds and insects in the Magdalena River Valley in Colombia (BAR radar), and in the diurnal birds and insects in the Tallaganda mountains in Southeast Australia (Fig. 3A-C). The peak bird proportions and bird volumes corresponded to both boreal and austral spring and fall in Colombia and Australia respectively (i.e. ∼ April and October). The reflectivity proportion of birds was similar in both countries, averaging around 0.48 ±0.17 (SD, reflectivity weighted) in Colombia and 0.53 ±0.19 in Southeast Australia. The north-south V component of the bird ground speed was mainly oriented northward in the movement peak around April and southward in the peak around October for both Colombia and Australia, and less concentrated during other times of the year (Fig. 3D), indicating a seasonal flip of migration direction in both sites consistent with the expected orientation of seasonal migration.

**Figure 3:**
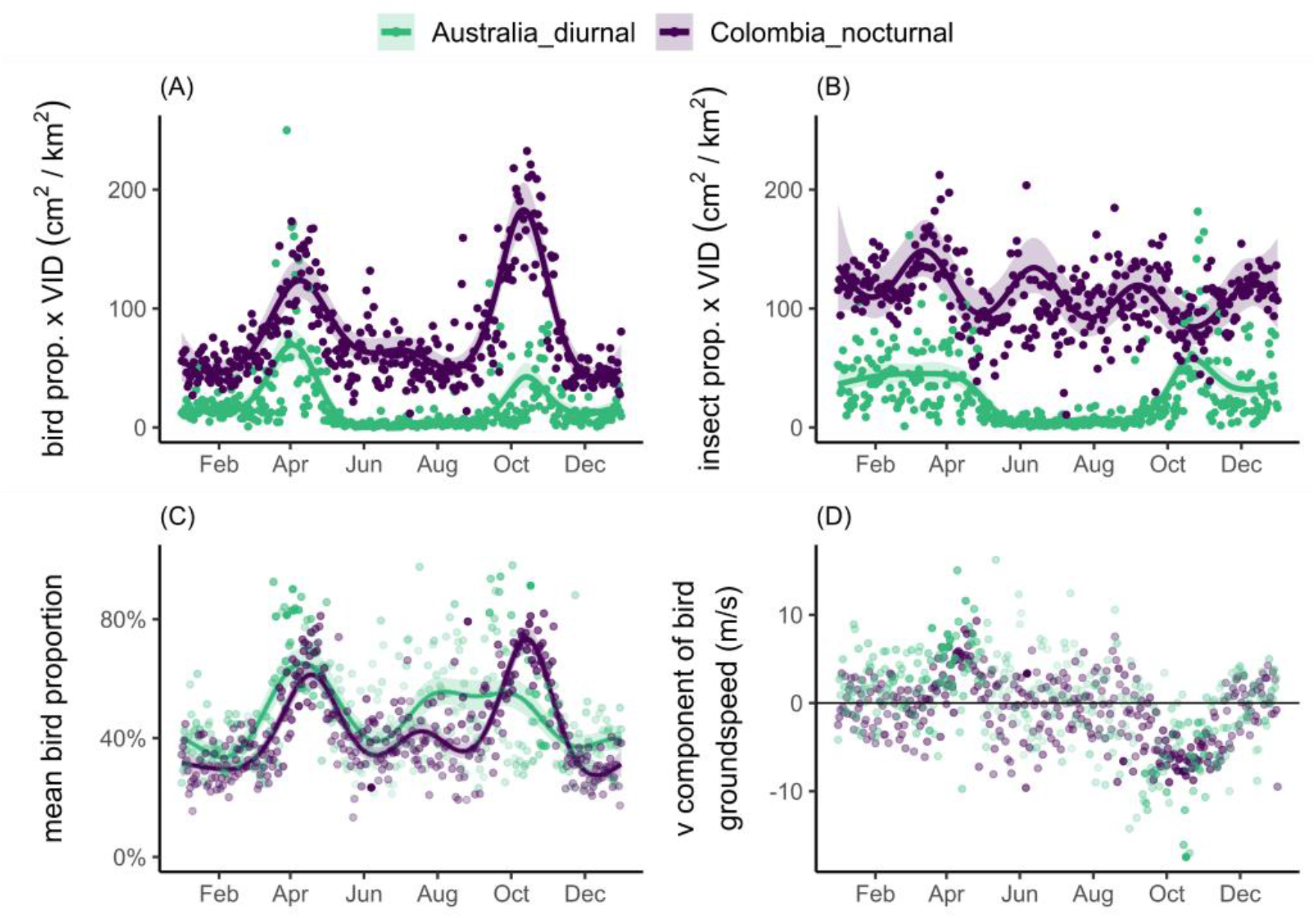
Seasonal variation in mean daily Vertically Integrated Density (VID, cm^2^/km^2^) for birds (A), insects (B), mean bird proportions for Australian and Colombian data (C) and v component of bird airspeed (D). Each point is a mean daily value of the focal parameter across multiple years of weather surveillance radar data at Captain’s Flat, Australia (*n* = 1 year) and Barrancabermeja, Colombia (*n =* 4 years). Solid lines and shaded 95% confidence bands are from GAM fit result. VID in Australia data are divided by 100 to facilitate comparison with Colombian data. Less transparency of the points indicating relatively larger VID in (C) and (D) and vice versa.

Seasonal peaks in Australian insect densities also correspond to non-migratory periods for birds, remaining high through late austral summer and becoming depressed during austral winter. In contrast, insect densities remained high throughout the year in Colombia, peaking at the beginning of the local rainy season around early April. Compared to filtering insects using a fixed airspeed criterion of 5 m/s, our method retained 8% and 11% more bird density in the Colombian data and Australian data respectively, and 63% and 78% of all time-altitude bins in the Colombian data and Australian data that would otherwise be removed using a 5 m/s threshold.

### Daily Patterns

Daily patterns of insect and bird dynamics reveals a multitude of patterns, for which we highlight one illustrative example for each study site. Across a 24-hour period during peak spring migration over the Guaviare radar in the Colombian Amazon, two discrete episodes of high reflectivity are apparent: one during daylight hours between 07:00 and 15:00 (local time) at elevations below 1000 m, and another that begins after sunset around 19:00 and continuing overnight through 04:00 (Fig. 4A). The nocturnal signal forms two layers, a weaker one below 1000 m and a stronger one around 3000 m, with a vertical band of stronger signal connecting them early in the night. This pattern matches the known phenomenon of nocturnally migrating birds taking off after sunset and ascending to altitudes with preferred wind conditions for flight (Dokter et al. 2013, Kemp et al. 2013, Horton et al. 2016b). Our approach evaluates this high-altitude nocturnal layer of reflectivity as entirely migrating birds (Fig. 4C). Visualizing the north-south (V) component of the wind (positive values are winds blowing from south to north and vice versa) further supports the hypothesis that this high-altitude reflectivity layer is birds as it corresponds with the altitude where birds reach minimally opposing winds above 2500 m for birds migrating north in the boreal spring (Fig. 4I). This is a known altitude selection strategy used by migrant songbirds (Dokter et al. 2013, Kemp et al. 2013). It is also unlikely that insects would be flying at high altitudes between 1000 and 3000 m in Colombia (Bell et al. 2013, Gao et al. 2024) In contrast, our model evaluates the reflectivity signal during the afternoon as mostly insects, when high temperatures and local convection offer ideal flight conditions for many insect taxa (Fig. 4G). We also see evidence of early morning bird migration after sunrise (Fig. 4C and E).

**Figure 4:**
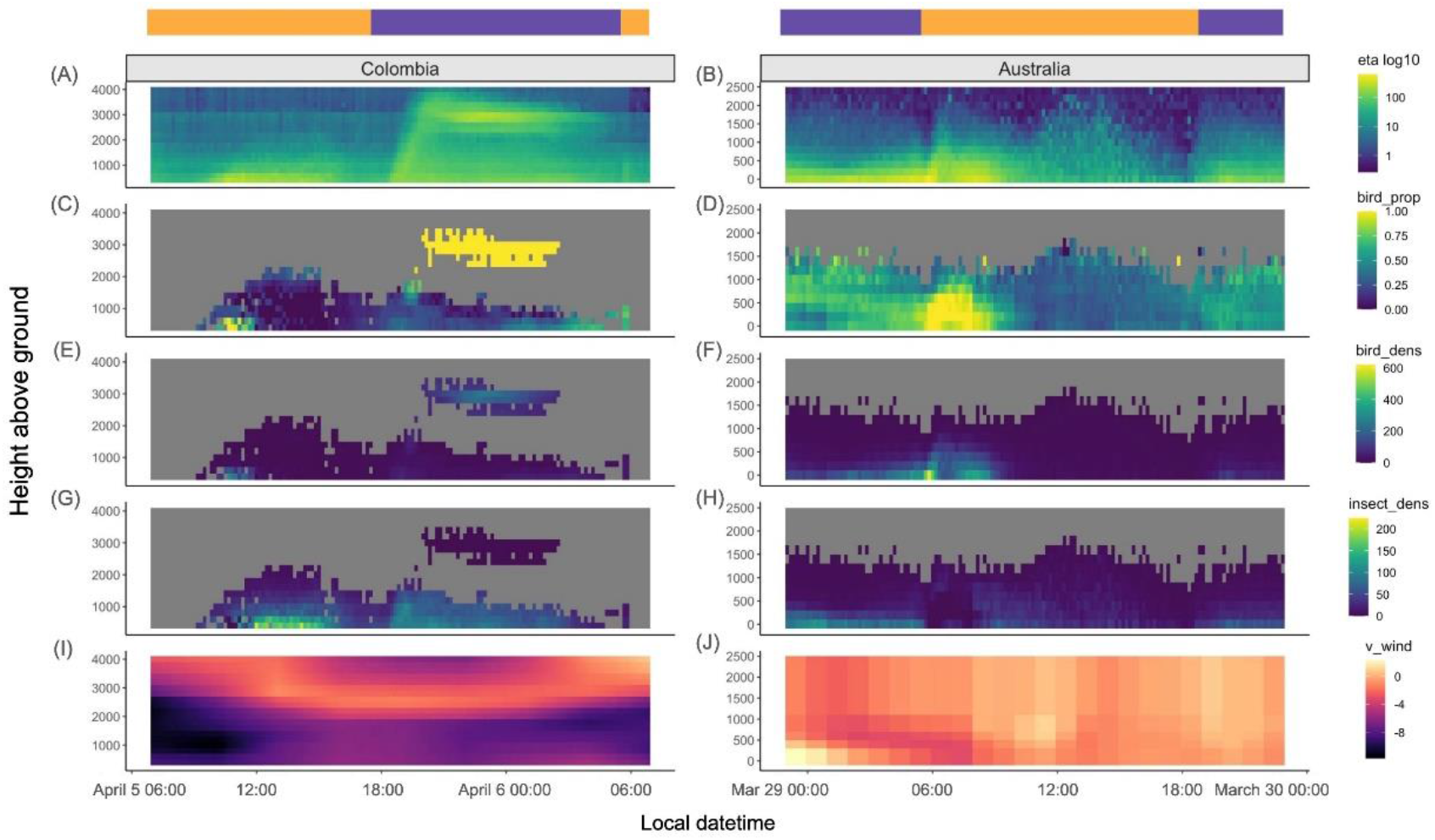
Variation in reflectivity (η, cm^2^·km^-3^), estimated bird proportions, estimated bird densities (bird proportion * η), estimated insect densities (1-bird proportion * η), and the north-south wind component (v_wind_) across example 24-hour periods in San Jose del Guaviare, Colombia in April 2019 and Captain’s Flat, Australia in March 2018. Colored bars above panels indicate whether the corresponding time on the x-axis is nocturnal (violet) or diurnal (orange). Gray areas lack sufficient velocity in the radar product to calculate bird proportions. Note different scales of y-axis for two sites.

Over Captains Flat in Southeast Australia, one distinct peak movement of high reflectivity occurred around the civil dawn (06:46) during the austral autumn in the twenty-four hour period of Mar 29, 2018, between 06:00 and 11:00 at elevations up to 1000 m above ground level (Fig. 4B). Such dawn peaks were commonly observed during March and April 2018, and to a lesser extent in Oct 2018, and not seen in winter and summer months. Weaker reflectivity persisted throughout the rest of the daytime and declined toward the sunset, and reflectivity increased after civil dusk (19:26) and persisted throughout the nocturnal period, mainly in the lower altitudinal bands. Our analysis indicates that the peak movement around dawn is almost entirely birds with high airspeed and thus high bird proportion (Fig. 4D). Bird proportion in the reflectivity declined sharply after 11:00 and remained low during the rest of the daytime, and increased once again after 20:00. We also find that insect activity shows a strong temporal contrast with bird activity (Fig. 4F and H). Insects dominated the daytime after 11:00 that can reach up to higher latitudes, showing a similar pattern as the Colombian case. Insect reflectivity persisted throughout the nocturnal period as well, mainly in the lowest altitudinal layer (Fig. 4H).

### Sensitivity Analysis

Simulated data showed that mean bird proportion, as calculated over the full dataset for each location, showed a negative relationship with increasing bird airspeed and a positive relationship with increasing insect airspeed (Fig. 5). The sensitivity of mean bird proportion differs with bird and insect airspeed: mean bird proportion increased from 0.42 to 0.51 when insect airspeed increased from 0 to 5 m/s but decreased from 0.42 to 0.23 when bird airspeed increased from 8 to 15 m/s using the Australian data. A similar pattern is observed in the Colombian data and overall, the mean bird proportion is more sensitive to the value of the bird airspeed parameter than the insect airspeed parameter.

**Figure 5.**
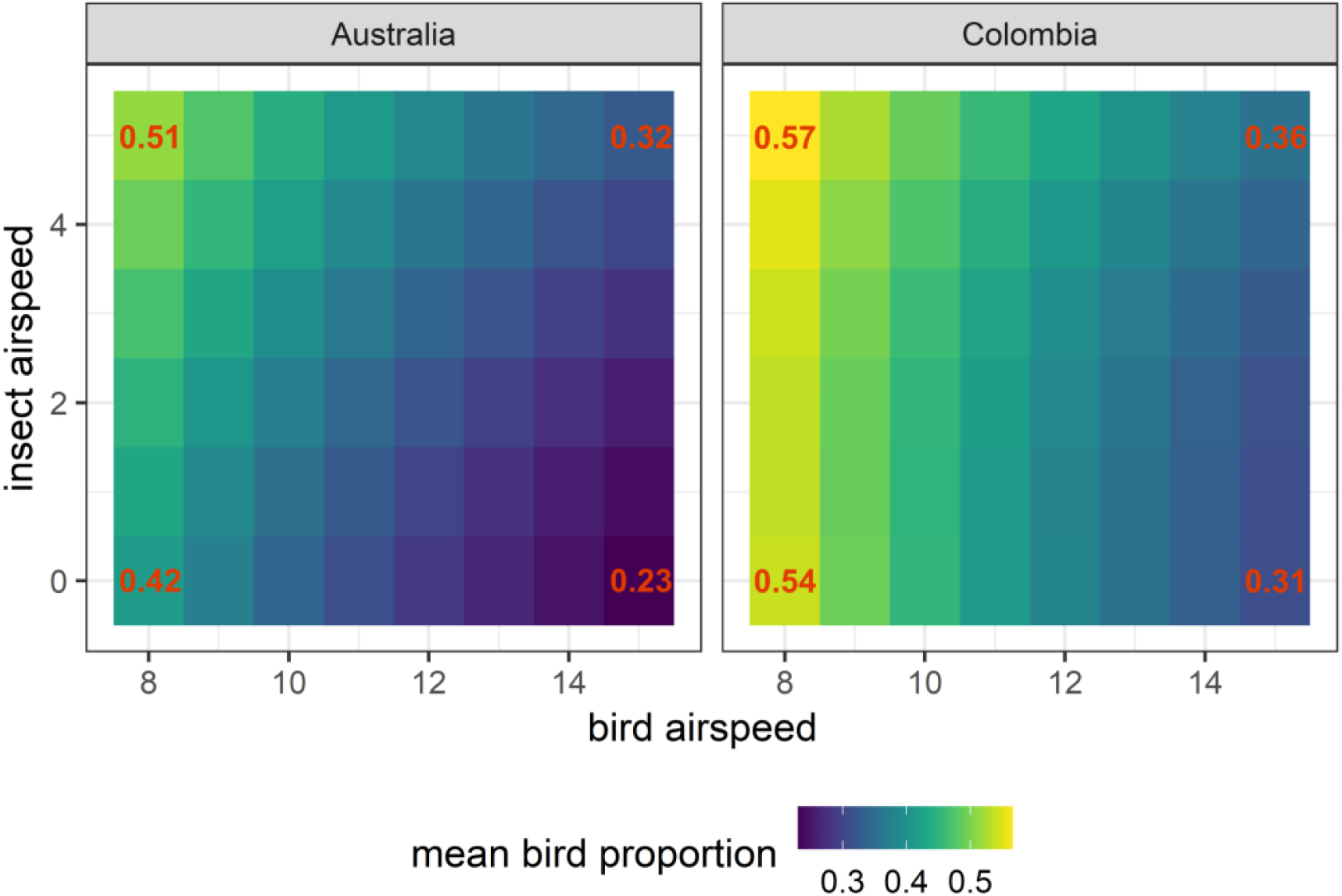
Evaluation of the sensitivity of retrieved mean bird proportion values as a function of the insect and bird airspeed parameters, using data from Barrancabermeja in Colombia and Captain’s Flat in Australia. Mean bird proportions at the four corners of each plot are labeled with the numerical values to assist interpretation. The color scale range covers values from 0.23 to 0.57 for visualization purposes, rather than the range of possible values between 0 and 1.

## Discussion

Our case studies illustrate that our approach is capable of recovering bird and insect signals that match expected biological patterns across daily and annual cycles in both Australia and Colombia. Bird migration activity increases during the expected periods of seasonal migration, and we find clear seasonal flips in migratory direction in the transition from spring to fall. Bird migration in Colombia is apparent especially at night, while in southeast Australia migration is predominantly diurnal, a characteristic that seems to be common among migratory Australian landbirds (Chan 2001). These seasonal patterns match the expected phenology of both birds and insects in their respective environments: in southern temperate regions, insect populations peak during the warmer months, which encompasses the main period of bird migration, while the activities of both taxa diminish during the austral winter. Therefore, migratory birds and insects co-occur in time more in temperate Australia, while birds emerge during migratory periods from a high baseline volume of aerial insects that is present year-round in tropical Colombia.

On a daily scale, we recover fine-scale dynamics in the altitudes and abundances of birds and insects. These detailed seasonal and daily patterns would not be detectable using traditional airspeed thresholding methods for separating birds and insects in weather radar data, which remain standard practice in the field (e.g. Cohen et al., 2021; Farnsworth et al., 2016; Horton et al., 2019, 2016b). Identifying taxa based on airspeed typically only works in non-mixture cases, where the airspace is dominated by either birds or insects at different times. However, in the locations of our case studies in tropical and south-temperate climates, birds and insects were found to co-occur at similar magnitudes in mixtures. As a result, the distribution of airspeed of all aerial biomass combined is often unimodal. Airspeeds are also weighted heavily towards lower values (Fig S1, Electronic Supplementary Material), even during heavy bird migration, because of the strong background of aerial insects. These results show that in these climates, separating bird and insect cases using a single airspeed threshold is problematic.

Using simple assumptions on the magnitude of the airspeed of birds and insects, our alternative approach is able to estimate proportions of birds and insects in our case studies under scenarios where it would not be possible using simply airspeed thresholding. Our model also allowed us to investigate fine-scale patterns in time and space of both birds and insects, by retaining a large proportion of time-altitude data points that would otherwise be excluded under fixed criteria. These data points are vital for providing a comprehensive depiction and revealing fine-scale structures of the time-altitude distributions of both taxa. Our approach is especially valuable for advancing aeroecology research in insect-rich areas like the tropics, parts of the southern hemisphere, and summer to early autumn periods in the northern temperate zone, when insects may outweigh birds in terms of reflectivity.

Our approach requires analysts to include explicit values for the expected bird and insect airspeeds. We argue that in most systems, reasonable estimates can be obtained from the expected species composition of migratory birds from the expected phenology of migration, combined with known self-powered speeds of individual species. Weather radar signals of birds tend to be heavily biased towards migratory passerines, because of their relatively high abundance compared to other species groups, especially at night (Dokter et al. 2018). Average passerine airspeeds detected by tracking radars tend to cluster in a fairly narrow range of values (10-14 m/s, Karlsson et al., 2012; Liechti, 1995; Nilsson et al., 2014), which vary with season, geography, or wind only to a limited extent (10-15%, Kemp et al., 2010; Nilsson et al., 2014). Notably, ground speeds (and conversely inferred airspeeds) detected by weather radars tend to be lower by approximately 1 m/s compared to those of tracked individuals (Nilsson et al. 2018), because weather radars detect speeds averaged over many individuals with some directional spread. Weather radars also have a much better ability to detect small insects that are more prone to full drift (Chapman et al. 2011, Hu et al. 2016), and meteorologists have used insects successfully as passive tracers of the wind field (Holleman 2005, Rennie 2014). We therefore expect insect airspeeds to be on the lower end of the possible range (∼1 m/s) to be appropriate in most cases. Given the inherent associated uncertainties in assumed airspeeds and available altitudinal wind data, we recommend testing the robustness of any conclusions against a range of assumed bird and insect airspeeds, or applying bootstrapping procedures around possible ranges of airspeeds to generate associated uncertainties in estimates.

Our sensitivity analysis indicates that inferred bird-insect proportions are much more sensitive to the assumed bird airspeed than insect airspeed, because bird airspeeds cover a potentially wider range of values. In our case studies we used fixed airspeeds throughout day and night and across the whole year. Importantly, airspeeds in both birds and insects may vary throughout the year. For example, at temperate latitudes early spring and late fall tends to be associated with migration of larger-bodied species than late spring and early fall (Dokter et al. 2011, Horton et al. 2019). With knowledge of local biology, that is, priors on which taxa are aloft during the time and place of interest, seasonally varying airspeed values can be easily implemented.

Nussbaumer et al. (2021b) showed that in a temperate climate the year-round aggregate bimodal distribution of airspeed and radial velocity standard deviations (sd_vvp), as obtained in a radial velocity fit using Volume Velocity Processing (vvp), may show distinct peaks that can be attributed to insect and bird components. Analysis of mixture airspeed distributions like this can be used to obtain data-driven estimates of bird and insect airspeeds to parametrize our equations. However, we have found that distinct bimodal distributions of airspeed are not always present, in which case one will need to rely on a priori or independent estimates. Also, we have found that S-band data has shown less clear variation in sd_vvp in S-band NEXRAD data compared to C-band data (Dokter pers. obs.), limiting the generality of the approach that relies on a bimodal distribution fit along both the sd_vvp and airspeed axes.

An advantage of our approach is its analytical simplicity combined with a requirement for analysts to be explicit about assumptions on self-powered airspeeds of birds and insects, respectively. As an indeterminate problem, these speeds cannot be measured directly by the radar in mixtures, and transparency about necessary inferences we believe benefits the reproducibility of results. Analytical simplicity also makes our approach well suited for applications that need to be fast and real-time, such as vertical profile extractions that feed operational archives and real-time observation systems like BirdCast (www.birdcast.info) and the European Aloft data repository (www.aloft.eu, Nilsson 2024, in review).

Importantly, due to the differences in scattering mechanism for birds and insects, the mixture proportions of insect and bird reflectivity depend strongly on the radar wavelength. In our study we found similar reflectivity proportions in Colombia and Australia. However, Australian data was collected at S-band and Colombian data at C-band. Due to the λ4 relationship with wavelength of insect’s radar cross section (Eq. 4), insects at S-band wavelengths (10 cm) show up a factor 24 = 16 times weaker than at C-band wavelengths (5 cm, twice as short). The insect component in Australia is therefore comparatively much stronger than in Colombia, all else being equal. The difference in the detected magnitude of the insect component also depends strongly on the insect size distribution (see Eq. 4), and could be either due to a higher number of insects or due to a distribution of larger sizes (this strong wavelength dependence is the main reason why meteorologists prefer to express reflectivity as Z instead of η, which removes the λ4 dependence for objects smaller than the radar wavelength, like hydrometeors and insects).

The computational ease of our model makes it effective for a variety of biological scenarios and data structures. However, imperative to its function is the presence of radial velocity data in the weather radar product that is used to calculate animal airspeeds, and the availability of high-quality wind data. While radial velocity is a standard measurement collected and stored by meteorological agencies around the world, it may be poorly represented in countries in the early stages of developing weather surveillance radar infrastructure (Shamoun-Baranes et al. 2022). For example, velocity data may be a second priority relative to reflectivity when data storage is less financially or logistically feasible, or operating protocols are in the process of being established. One common symptom of weak velocity data is poor information when reflectivity is low, driving much of the gray areas in our vertical profiles from Colombia. We urge meteorological agencies to support velocity data in their radar products to the extent possible.

## Conclusion

Distinguishing birds and insects has been a central challenge in the field of aeroecology. We demonstrated that by incorporating simple and explicit assumptions on the self-powered airspeeds of birds and insects, we can achieve separation of their contributions to volumes of aerial biomass in weather radar data and produce ecologically meaningful patterns of both taxa across diverse climates. By accurately identifying bird migration patterns, it can help pinpoint critical habitats and migratory corridors that need protection, and assist in mitigating risks associated with bird collisions with man-made structures such as buildings and wind energy structures, by providing detailed migration timing and routes. This new tool to infer the broad taxonomic composition of aerial biomass invites application across many research questions, study systems, and spatial-temporal scales. Our analytical expressions can be easily implemented into existing software packages and readily applied on vertical profiles of radar data. These features can empower ecologists to efficiently analyze millions of previously unexplored radar scans from regions other than the northern temperate zone, as well as diurnal migration of birds and insects that are comparatively understudied.

