## Supplementary materials for "Analysis of mixtures of birds and insects in weather radar data"

### Supplementary methods

For meaningful ecological assumptions, we would have $a_{bird}>a_{mix}>a_{insect}$. Therefore both $p_{1}$ and $p_{3}$ < 0. $\sqrt{p_{2}^{2}-4p_{1}p_{3}}$ would be always smaller than the absolute value of $p_{2}$, and $p_{2}$ must be positive for $f$ to be above zero in (22).

Given that $w^{2}={||\boldsymbol{w}||}^{2}$and assuming $\theta$ is the angle between $\boldsymbol{g}_{mix}$ and $\boldsymbol{w}$, $p_{2}$ can be rewritten as:

$$\begin{aligned} p_{2}={2a_{bird}}^{2}-2*\left( \frac{a_{insect}}{w} \right)*\left( \boldsymbol{w}\cdot\boldsymbol{a}_{mix} \right)=2*\left( {a_{bird}}^{2}-a_{mix}*a_{insect}*cos \theta\right) \#\left( 15 \right) \end{aligned}$$

Since $a_{bird}>a_{mix}>a_{insect}$, $p_{2}$ will always be positive.

Therefore, in (13) the solution that adds the square root would always be smaller than the solution that subtracts the square root, and more likely to be a valid proportion between 0 and 1. In the solution that subtracts, for $f$ to be a valid proportion that is smaller than 1 (and given that $p_{1}$ is negative), it would need:

$$\begin{aligned} {-p}_{2}-\sqrt{p_{2}^{2}-4p_{1}p_{3}}>2p_{1} \#\left( 16 \right) \end{aligned}$$

Or:

$$\begin{aligned} p_{2}^{2}-4p_{1}p_{3}<\left( p_{2}+2p_{1} \right)^{2} \#\left( 17 \right) \end{aligned}$$

Simplify the above equation, we have:

$$\begin{aligned} p_{3}{+ p}_{2}+p_{1}<0 \#\left( 18 \right) \end{aligned}$$

Combing (10), (12), (15) and (18), we have:

$$\begin{aligned} {a_{insect}}^{2}-{a_{bird}}^{2}+2*\left( {a_{bird}}^{2}-a_{mix}*a_{insect}*cos \theta\right)+{a_{mix}}^{2}-{a_{bird}}^{2}<0 \#\left( 19 \right) \end{aligned}$$

The left side of (19) would be smallest if $cos \theta=1$. In this condition, the above equation can be simplified as:

$$\begin{aligned} \left( b-{ff}_{a} \right)^{2}<0 \#\left( 20 \right) \end{aligned}$$

Which is not true. Hence $f$ has only one solution:

$$\begin{aligned} f=\frac{-p_{2}+\sqrt{p_{2}^{2}-4p_{1}p_{3}}}{2p_{1}} \#\left( 21 \right) \end{aligned}$$

And we can subsequently use equation (11) and (12) to solve $u$ and $v$, and equation (10) to solve the heading of the birds.

$f$ will have real number solution if:

$$\begin{aligned} p_{2}^{2}-4p_{1}p_{3}>0 \#\left( 22 \right) \end{aligned}$$

combing (10), (12), (15) and (22), we have:

$$\begin{aligned} \left( 2*\left( {a_{bird}}^{2}-a_{mix}*a_{insect}*cos \theta\right) \right)^{2}>4*\left( {a_{insect}}^{2}-{a_{bird}}^{2} \right)*\left( {a_{mix}}^{2}-{a_{bird}}^{2} \right) \#\left( 23 \right) \end{aligned}$$

if $cos \theta<0$, the left side of (23) will always be larger than $2*{a_{bird}}^{2}$, and the smallest possibility is when $cos \theta\to0,$then (23) will become:

$$\begin{aligned} \left( {a_{bird}}^{2} \right)^{2}>\left( {a_{insect}}^{2}-{a_{bird}}^{2} \right)*\left( {a_{mix}}^{2}-{a_{bird}}^{2} \right) \#\left( 24 \right) \end{aligned}$$

which is always true. If $cos \theta>0,$the left side of (23) will always be larger than zero, and the smallest possibility is when $cos \theta=1$, which will become:

$$\begin{aligned} \left( {a_{bird}}^{2}-a_{mix}*a_{insect} \right)^{2}>\left( {a_{insect}}^{2}-{a_{bird}}^{2} \right)*\left( {a_{mix}}^{2}-{a_{bird}}^{2} \right) \#\left( 25 \right) \end{aligned}$$

or

$$\begin{aligned} \left( a_{mix}-a_{insect} \right)^{2}>0 \#\left( 26 \right) \end{aligned}$$

Which is always true, therefore (22) is always true and (21) always has a real number solution.

### Supplementary results

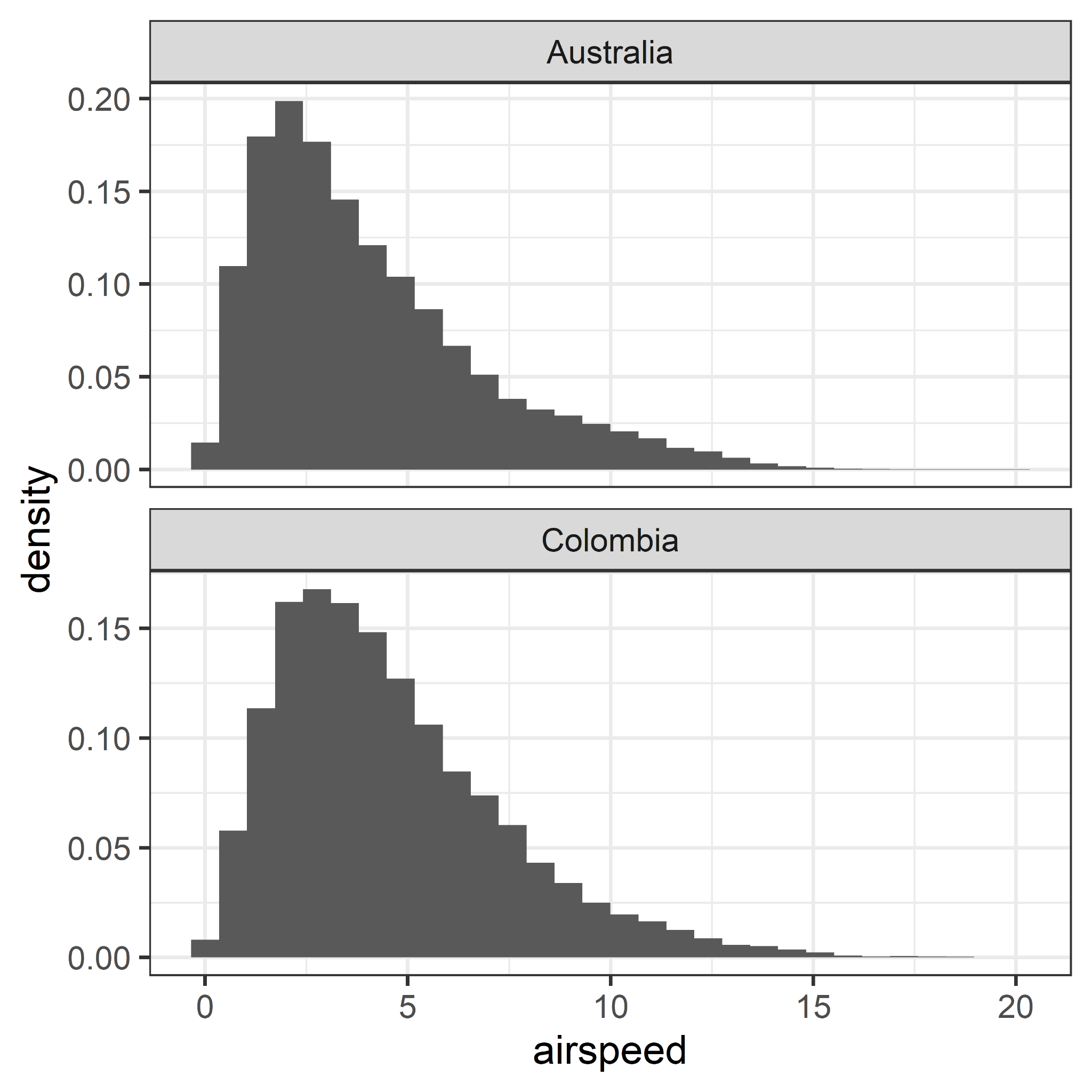

**Figure S1.** Distribution of mixture’s airspeed weighted by reflectivity in data from Barrancabermeja in Colombia and Captain's Flat in Australia. The airspeeds above 20 m/s are removed from plotting.

### Supplementary codes:

\# R code for calculating insect proportion f and bird proportion 1-f

#' Partition mixtures of animals using assumptions on airspeeds.

#'

#' Partition mixtures of animals using assumptions on airspeeds.

#' @param x a mixture animal density or linear reflectivity eta.

#' @param u the mixture's ground speed u component (west to east)

#' @param v the mixture's ground speed v component (south to north)

#' @param U the west to east wind component

#' @param V the south to north wind component

#' @param fast the fast component's airspeed

#' @param slow the slow component's airspeed

#' @param drop_slow_component when TRUE (default) output density, ground speed and

#' heading for fast component, when FALSE for slow component.

#' @return a data.frame with corrected density or reflectivity x, ground speed (u, v)

#' and heading in clockwise degrees from north.

#' @export

#' @examples

#' # drop the slow component (typically insects)

#' clean_mixture(100,-13,13,-7,6, fast=12, slow=1)

#' # drop the fast component (typically birds)

#' clean_mixture(100,-13,13,-7,6, fast=12, slow=1, drop_slow_component=FALSE)

clean_mixture <- function(x, u, v, U, V, slow = 1, fast = 12, drop_slow_component = TRUE){

\# verify input

assertthat::assert_that(x >= 0)

assertthat::assert_that(assertthat::is.number(u))

assertthat::assert_that(assertthat::is.number(v))

assertthat::assert_that(assertthat::is.number(U))

assertthat::assert_that(assertthat::is.number(V))

assertthat::assert_that(slow >= 0)

assertthat::assert_that(fast > 0)

assertthat::assert_that(fast > slow)

assertthat::assert_that(is.flag(drop_slow_component))

\# define helper quantities:

wind_speed <- sqrt(U^2 + V^2)

wind_direction <- atan2(V,U)

mixture_airspeed <- sqrt((u-U)^2 + (v-V)^2)

\# catch limiting cases

if(mixture_airspeed > fast){

warning("Airspeed of mixture exceeds airspeed of fast component, assigning all weight to fast component")

f=0

}

if(mixture_airspeed < slow){

warning("Airspeed of slow component exceeds airspeed of mixture, assigning all weight to slow component")

f=1

}

if(mixture_airspeed <= fast & mixture_airspeed >= slow){

p1 <- slow^2 - fast^2

p2 <- 2*fast^2 - 2*(slow/wind_speed)*(u*U+v*V - wind_speed^2)

p3 <- (u-U)^2 + (v-V)^2 - fast^2

\# signal proportion attributed to the slow component:

f <- (-p2+sqrt(p2^2-4*p1*p3))/(2*p1)

}

if(drop_slow_component){

\# fast component airspeed, typically birds:

if(f==1){

air_u=NA

air_v=NA

} else{

air_u=((u-U)-(slow/wind_speed)*U*f)/(1-f)

air_v=((v-V)-(slow/wind_speed)*V*f)/(1-f)

}

x_corr=(1-f)*x

} else{

\# slow component airspeed, typically insects:

air_u=slow*cos(wind_direction)

air_v=slow*sin(wind_direction)

x_corr=(f*x)

}

data.frame(x=x_corr,u=U+air_u,v=V+air_v,heading=(pi/2atan2(air_v,air_u))*180/pi)

}
